# Enrichment of Zα domains at cytoplasmic stress granules is due to their innate ability to bind nucleic acids

**DOI:** 10.1101/2021.01.20.427402

**Authors:** Luisa Gabriel, Bharath Srinivasan, Krzysztof Kuś, João F. Mata, Maria João Amorim, Lars E.T. Jansen, Alekos Athanasiadis

## Abstract

Zα domains are a subfamily of winged Helix-Turn-Helix (wHTH) domains found exclusively in proteins involved in the nucleic acids sensory pathway of vertebrate innate immune system and host evasion by viral pathogens. Interestingly, they are the only known protein domains that recognise the left-handed helical conformation of both dsDNA and dsRNA, known as Z-DNA and Z-RNA. Previously, it has been demonstrated that ADAR1 and ZBP1, two proteins possessing the Zα domains, localize to cytosolic stress granules. It was further speculated that such localization is principally mediated by Zα domains. To characterize and better understand such distinct and specific localization, we characterised the *in vivo* interactions and localization pattern for the amino terminal region of human DAI harbouring two Zα domains (Z_αβ_^DAI^). Using immunoprecipitation and mass spectrometry, we identified several interacting partners that were components of the complex formed by Zα domains and RNAs. Differential interacting partners to wild-type Zα, relative to mutant proteins, demonstrated that most of the physiologically relevant interactions are mediated by the nucleic acid binding ability of the Z_αβ_. Further, we also show enrichment of selected complex components in cytoplasmic stress granules under conditions of stress. This ability is mostly lost in the mutants of Z_αβ_^DAI^ (Z_αβ_^DAI^ 4×mut) that lack nucleic-acid binding ability. Thus, we posit that the mechanism for the translocation of Zα domain-containing proteins to stress granules is mainly mediated by the nucleic acid binding ability of their Zα domains. Finally, we demonstrate that FUS and PSF/p54nrb, two RNA binding proteins with established roles in stress granules, interact with Zα, which provides strong evidence for a role of these proteins in the innate immune system.

## Introduction

Zα domains belong to an an unusual nucleic acids binding domain family which recognises purine/pyrimidine repeats (Kim et al., 2004; Oh et al., 2002) in the left-handed conformation of both DNA and RNA helices (Placido et al., 2007; Schwartz et al., 1999), both *in vitro* and *in vivo*. The prototypic Zα domain was found as part of large isoform of ADAR1, an interferon (IFN) inducible RNA editing enzyme, and all subsequently identified Zα domains were found to be components of IFN-inducible proteins or viral inhibitors of the interferon pathway (Athanasiadis, 2012). ADAR1 is shown to be a negative regulator of the dsRNA sensing pathway and has an important role in preventing activation of the pathway by self-RNAs. Mutations in ADAR1, some within the Zα domain, have been implicated in causation of the Aicardi-Goutières syndrome, an auto-inflammatory genetic disease (Rice et al., 2012). The other two cellular proteins that have Zα domains are DAI and PKZ. DAI is a mammalian sensor of dsDNA (Takaoka et al., 2007) while PKZ is a fish-specific effector of dsRNA recognition, having a role analogous to PKR (Rothenburg et al., 2005). Two more proteins containing Zα domains are both encoded by DNA viruses: E3L from poxviruses (Kim et al., 2003) and ORF112, a protein we recently described in a subfamily of Herpes viruses (Kus̈ et al., 2015). Both viral proteins have a role in innate immune evasion and are required for viral proliferation. In the case of E3L, it was shown that while a virus with deleted E3L Zα domain loses pathogenicity, replacement of the domain with Zα domains from either ADAR1 or DAI fully restores pathogenicity (Kim et al., 2003) suggesting that Zα domains from different proteins have similar function.

While the degree of conservation among Zα domains is moderate, key DNA/RNA interacting residues, identified in crystal structures of Zα/DNA/RNA complexes (Placido et al., 2007; Schwartz et al., 1999), are absolutely conserved and thus provide a motif for the unambiguous identification of the domains which otherwise demonstrate the common wHTH fold (Athanasiadis, 2012). Studies of Zα domains have primarily focused on the mechanism of their interaction with Z-DNA and Z-RNA through mostly structural and biochemical studies (De Rosa et al., 2010; De Rosa et al., 2013; Kim et al., 2011; Schwartz et al., 2001; Sung et al., 2008). Thus, although now there are structures of Zα domains in complexes with nucleic acids from all known Zα containing proteins (De Rosa et al., 2013; Ha et al., 2004; Kus̈ et al., 2015; Schwartz et al., 1999; Schwartz et al., 2001), little is known regarding the *in vivo* target(s) and function of the domains.

Zα domains *in vitro* can bind the double helix of both DNA and RNA taking advantage of the fact that, unlike the right-handed helices of the two macromolecules that have very different structures (B and A respectively), their left-handed helix is very similar (Placido et al., 2007). Thus the question whether Zα domains target DNA or RNA *in vivo* is open. Nevertheless, the fact that proteins containing Zα domains are predominantly cytoplasmic and have a function in the RNA sensing pathway strongly points to a Zα association with dsRNA and/or RNA/DNA hybrids. An affinity of Zα domains for segments of ribosomal RNA (Feng et al., 2011) has been shown through analysis of RNA pulldowns in both *E. coli* and human cell extracts. The same study showed that that expression of Zα domains can inhibit translation, although it is not clear if this inhibition is the result of binding to the suggested ribosomal RNA sites. In another work investigating the Z-DNA content of the genome with Zα domains as reporter, it was reported that Zα can interact with centromeric repeats (Li et al., 2009). In both cases, the physiological relevance of the described interactions remains unexplored and no clear link is established for the function of Zα containing proteins.

Two of the Zα containing proteins ADAR1 and DAI, have been shown to associate with stress granules (Deigendesch et al., 2006; Weissbach and Scadden, 2012). Stress granules are cytoplasmic RNPs containing stalled ribosomes, the result of stress-induced translation arrest. It has been shown that hyperedited dsRNA (IU-dsRNA) specifically accumulates in a stress granule like complex and suppresses interferon induction and apoptosis (Scadden, 2007; Vitali and Scadden, 2010). Further, it has been previously demonstrated that IU-dsRNA, apart from its role in interacting with protein-complex that largely comprised components of cytoplasmic stress granules, also interacted with proteins that are not known to have a role in stress granule formation (Callebaut and Mornon, 1997; Scadden, 2005; Yang et al., 2007). Recent work shows that Zα domains in isolation associate with SGs and can drive proteins to localise there (Ng et al., 2013). These results suggest that Zα domains may be responsible for the association of ADAR1 and DAI in SGs. Nevertheless, the mechanism of SG association and the interactions of Zα domains and other constituents within SGs are unknown.

Here we use the N-terminal part of human DAI, which consists of two Zα domains, to study its localisation and characterize the complexes in which they participate.

## Materials and Methods

### Cloning and Mutagenesis

The amino-terminal region of human DAI (hZαβ^DAI^, residues Ala2-Tyr165) was amplified by PCR using a construct of human GST-DAI (originating from NM_030776.2) as a template and primers incorporating XhoI and BamHI restriction sites (#1 and #2 in Table S1). The digested PCR product was cloned into a pIC113 vector which allowed the insertion of a LAP-tag (Cheeseman and Desai, 2005; Li, 2011) at the N-terminus providing fused Green fluorescent protein (GFP) and an S-tag with a cleavable TEV site between them (plasmid pIC113hZ_αβ_^DAI^). The entire construct or the LAP-tag only (to be used as control) was subcloned into pBABE^puro^ using primers (#3-5) and BamHI and SalI restriction sites yielding plasmids pBABEZ_αβ_^DAI^ and pBABE*GFP*. Site-directed mutagenesis was performed using the NZYMutagenesis kit (NZYTech), to mutate four hZαβ^DAI^ residues known to be critical for nucleic acid binding to Ala (Z_αβ_-N46A/Y50A and Z_αβ_-N141A/Y145A). The mutant constructs, pIC113hZ_αβ_^DAI^4xMut and pBABEZ_αβ_^DAI^4xMut, were generated following the manufacturer’s protocol with primers #6-9. The quadruple mutant would be called Z_αβ_^DAI^4xMut throughout the paper.

### Cell culture and arsenite treatment

A549 cells (human lung adenocarcinoma) and HEK293-G cells and their derivatives were cultured in Dulbecco’s Modified Eagle Medium (DMEM) supplemented with 10 % heat inactivated fetal bovine serum, penicillin (100 U per ml), streptomycin (100 μg per mL) and L-glutamine (2 mM) (complete media) in a 5 % CO_2_ humidified incubator at 37 °C.

To induce stress granule formation, A549 cells and their derivatives were cultured in Dulbecco’s Modified Eagle Medium (DMEM) supplemented with sodium arsenite (0.5 mM) for 30 min in 5% CO_2_. Before and after incubation with arsenite, cells were washed 3 × with PBS. Cells were then left to recover for 30 min in complete media lacking arsenite.

### Transient transfection and generation of stable cell lines

A549 cells were transfected with pIC113hZαβ^DAI^/pIC113hZαβ^DAI^4xMut using Lipofectamine LTX, PLUS reagent and OptiMEM (Life Technologies) according to manufactures instructions.

For the generation of stably expressing cell lines, HEK293-GP cells were co-transfected with pBABEZαβ^DAI^/pBABEZαβ^DAI^4xMut/pBABE^GFP^ and pVSV-G using Lipofectamine LTX, PLUS reagent and OptiMEM (Life Technologies), according to the manufacturer’s recommended protocol (Morgenstern and Land, 1990). Retroviral particles were collected after 3 days, filtered with 0.45 μm filters (Acrodisc, Pall) and stored at −80 °C. These were used to transduce A549 cells in the presence of 8 μg/mL of hexadimethrine bromide (Polybrene, Sigma). Following 24 hours of transduction, cells were trypsinized and seeded for selection in complete media supplemented with 4 μg/mL puromycin (Calbiochem, Merck Millipore). All cell lines were passaged 3 times before use and kept in culture supplemented with 2 μg/mL puromycin and/or stored in liquid nitrogen.

### Immunofluorescent staining and confocal microscopy

The following antibodies were used in this study: Anti-ZBP1 clone RonoB19 (WB 1:500, ref. 651602, Rat, BioLegend), anti-TIAR (C-18) (IFA 1:500, ref. SC-1749, Goat, Santa Cruz), anti-NONO (IFA and WB 1:500, ref. N8664, Rabbit, Sigma); anti-FUS (IFA 1:100, WB 1:500, ref. HPA008784, Rabbit, Sigma), anti-PSPC1 (N-terminal) (IFA 1:250 Ref. SAB4200068, Rabbit, Sigma), anti-PML (IFA 1:100, ab53773, Rabbit, Abcam) and anti-Coilin (IFA 1:100) (Rabbit, a kind gift from Dr. P. Navarro-Costa) all against the human proteins. Secondary fluorophore-coupled antibodies were all from the Alexa Fluor range (568 ref. A10042 and 647 ref. A21469, IFA 1:1000, Life Technologies).

Cells were fixed in 4 % formaldehyde solution in PBS for 20 min at room temperature (RT). After washing with PBS, samples were permeabilized with 0.2 % 100 × Triton (v/v) solution in PBS and 1 % fetal bovine serum at RT for 7 min. Cells were blocked and all subsequent immuno-fluorescence staining (incubation and washes) were performed in PBS containing 1 % fetal bovine serum. Cells were then incubated with the corresponding primary antibodies for 1 hour at RT or overnight at 4 °C (Coilin and PML). After 3 washes, samples were incubated with the respective secondary antibodies and DAPI (Gerbu BiotechniK) for 30 min at RT. Cells were then mounted using Fluoromount-G (eBioscience) mounting media.

Confocal images were acquired on a Leica SP5 confocal microscope using a 63 × 1.3NA oil immersion objective employing 405 nm, 488 nm, 568 nm and 633 nm laser lines. Spectral detection adjustments for the emission of DAPI, eGFP, Alexa 568 and Alexa 633 fluorochromes were made using HyD detectors in standard mode.

Z-stacks were acquired on a Leica high content screening microscope, based on Leica DMI6000 equipped with a Hamamatsu Flash 4.0 LT sCMOS camera, using a 100 × 1.44 NA objective, DAPI, eGFP, TRITC and Cy5 fluorescence filter sets and controlled with the Leica LAS X software. Images were processed using ImageJ software (NIH).

### Pulldown and Western blotting

A549 cells with or without arsenite treatment from 20, 15 cm confluent dishes were lysed in lysis buffer (10 mM Tris pH 7.4, 150 mM NaCl, 0.5 mM EDTA and 0.5 % NP40) complemented with 1 × protease inhibitors (Pierce Protease Inhibitor Mini Tablets, EDTA-Free, Thermo Scientific) and 2 U Turbo DNase (Ambion) on ice for 1 hour. The resultant cell lysate was centrifuged at 17000 rpm for 20 min at 4 °C. Further, the supernatant from the previous round was centrifuged a second time under the same conditions. The lysate was diluted to a total volume of 9 mL with dilution/wash buffer (lysis buffer with 0.05 % Tween20 instead of NP40) supplemented with protease inhibitors. 100 μl of GFP-trap magnetic beads (ChromoTek) were added to the suspension and incubated for 90 min on a roller device. Beads were magnetically separated using a magnetic stand (Merck Millipore) and washed 5 times with 1 mL ice-cold dilution/wash buffer. The proteins of interest were eluted by cleavage with 10 units of AcTEV protease (Life Technologies) in a buffer containing 50 mM Tris HCl pH 8.0, 0.5 mM EDTA and 1 mM DTT in a total volume of 60 μL. The mix was incubated for 90 min on ice. For RNAse treated samples, the supernatant obtained after centrifugation of cell lysate was split in two and then incubated with beads in the presence of either RNAse cocktail (Ambion) 400 U of RNAse T1 and 10 U RNAse or 400 U RNAse inhibitor (SuperRNAseIn, Invitrogen). The rest of procedure was similar to that employed for RNAse treatment except that the dilution/wash buffer contained 5 mM MgCl_2_. Final samples were stored at −80 °C. For the western blot, samples were prepared after addition of 2.5 % β-mercaptoethanol to sample buffer (NuPAGE LDS sample Buffer, Life Technologies) and boiling for 5 min at 90 °C. The samples were electrophoresed on NuPage 4-12 % Bis-Tris SDS precast gels from Invitrogen. In all cases, Novex sharp pre-stained molecular weight marker (Life Technologies) was loaded for sizing purposes.

Polyacrylamide gels were then blotted on iBlot Gel Transfer Stacks (Nitrocellulose Mini) following manufacturer’s instructions for iBlot transfer device (Invitrogen). Membranes were blocked in blocking buffer (PBS 0.1 % Tween20 and 5 % milk powder) for 30 min at RT. After 3 washes (PBS 0.1 % Tween), membranes were incubated with corresponding antibodies for 60 min at RT. Then membranes were washed 3 × with (PBS 0.1 % Tween20) and incubated for 30 min at RT with the corresponding secondary antibodies IRDye 680RD (Ref. 926-68073, Li-Cor) or DyLight 800 (Ref. 039612-145-120, Rockland) (1:10000 dilution). Subsequently, the membranes were washed 3 times in buffer containing PBS with 0.1 % Tween20 and 2 times in PBS alone. Western blots were imaged using a LI-COR Biosciences Odyssey near-infrared scanner.

## Mass spectrometry for identification of the proteins

The mass spectrometry was performed at UPF-CRG proteomics facility (Center of Genomics Regulation (CRG)), Barcelona, Spain. Liquid chromatography coupled tandem Mass spectrometry (LC MS/MS) was carried out with LTQ-Orbitrap (Thermofisher scientific). Trypsin and LysC, a serine endopeptidase, was used in solution to carry our partial digest of the pull-down. The resulting digested peptide-containing solution was desalted by reverse phase and the peptides were fractionated with a C18 column. Fragmentation was carried out with in-source collision induced decay (Is CID) (collision energy 0-1000). Ionization source was in-source nanospray and the mass-spec was carried out in positive ion mode.

### Raw Data Processing

All raw files were analyzed together using the software MASCOT. Frequently observed contaminating proteins as reported in Crapome (Mellacheruvu et al., 2013) such as Myosin, Tropomyosin, Keratin, Actin, Trypsin etc, though reported diligently, were not given any weightage in further analysis. Carbamidomethylation of cysteine was set as a fixed modification (57.021464 Da) and N-acetylation of proteins N termini (42.010565 Da) and oxidation of methionine (15.994915 Da) were set as variable modifications. As no labeling was performed, multiplicity was set to 1. During the main search, parent masses were allowed an initial mass deviation of 7 ppm and fragment ion’s mass tolerance was set to 0.5 Da. PSM and protein identifications were filtered using a target-decoy approach at a stringent false discovery rate (FDR) of 0.01 % and a relaxed FDR of 0.05 %.

### Post-Processing

The final score was computed based on the peptide score match (PSM). The final score obtained for each experiment (e.g. Z_αβ_^DAI^ + arsenite, Z_αβ_^DAI^ – arsenite, Z_αβ_^DAI^ 4×mut + arsenite, Z_αβ_^DAI^ 4×mut – arsenite and GFP +arsenite) were arranged in experimental triplicates and the mean and standard deviation of each row, representing values obtained for each unique interacting protein, was computed. Student’s *t*-test was performed comparing the bait pull-down (in triplicates) to its individual bait specific control group (in triplicates). This control group varied depending on the nature of comparison i.e., proteins (mutant and wild type) vis-a-vis GFP, mutant vis-a-vis wild-type and proteins (mutants and wild-type) untreated vis-à-vis treated. This whole procedure of individual clustering, statistical parameter computation and *t* test was repeated for every bait. The resulting differences between the log_2_ means of the two groups (“log2(bait/background”) and the negative log *p*-values were plotted as volcano plots using GraphPad Prism ver 7. A stringent cut-off of 4 was set for log_2_ fold-change representing at least 16-fold change of the test with respect to the control group. A value of 2 for –log(*P*-value), corresponding to a *P*-value of <0.01, was employed to ascribe statistical significance. Quadrants are employed for graphical clarity in Figs and Tables explicitly state whether the imputed scores are significant statistically (P-value), numerically (fold-change) or additive (both statistical and numerical).

## Results

### The Z_αβ_^DAI^ domain localizes to stress granules

ADAR1 and DAI, the two mammalian proteins that contain Zα domains, localise to cytoplasmic RNPs known as stress granules (SGs) (Deigendesch et al., 2006; Weissbach and Scadden, 2012). Previous work from us and others has suggested that isolated Zα domains also localise to stress granules (Kus̈ et al., 2015; Ng et al., 2013) and thus may be responsible for the localisation of the proteins that contain them. To further explore the specific subcellular distribution of Zα domains, their nucleic acid targets and potential protein interactors, we constructed A549 cells stably expressing a GFP fusion to the N-terminal 164 aa of the nucleic acids sensor DAI which contains two Zα domains. We expressed this GFP-Z_αβ_^DAI^ fusion in stably transfected A549 cells and determined the distribution of the protein under different growth conditions. Under unperturbed conditions, the GFP-Z_αβ_^DAI^ fusion shows a diffuse distribution in both the cytoplasm and nucleus (Fig 1A bottom panel). When cells are treated with arsenite, Z_αβ_^DAI^ becomes enriched in cytoplasmic granules (Fig 1A top panel) that we identified as SGs by TIAR staining (Fig 1A top panel). In addition, we observe intense staining to a variable number of nuclear foci that resemble paraspeckles in size, shape and distribution (Fig S1) (For details refer to, Supplementary Information results). However, these nuclear structures appear only after arsenite treatment suggesting they are components of stress responses. As both SGs and paraspeckles are RNPs, these results suggest that Zα domains may be binding to RNAs that localise in such bodies. To understand whether Z_αβ_^DAI^ has a specific role in stress granule physiology and localizes to stress granules specifically or alternatively due to a secondary effect because of its affinity for ribonucleic acid, we decided to characterise the components of Zα complexes. Immunoprecipitation (IP) was performed employing anti-GFP beads and the resulting IPs were then used to identify proteins involved in Z_αβ_^DAI^ complexes.

**Figure 1.**
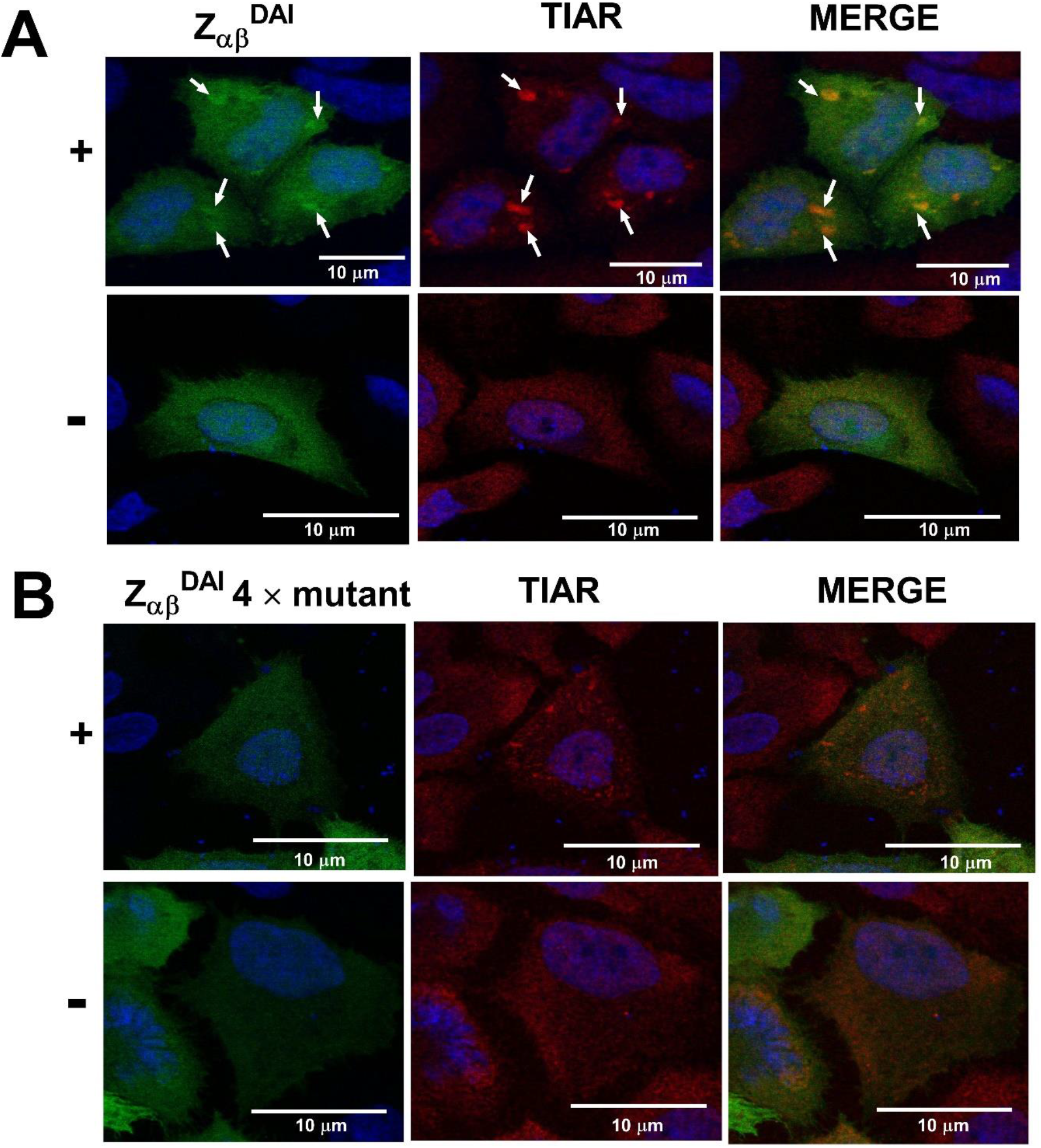
Z_αβ_^DAI^ is enriched in stress granules. **(A)** Immunolocalization of GFP-Z_αβ_^DAI^ fusion (green) in the presence (+) and absence (−) of stress induced by arsenite in A549 cells shows that GFP-Z_αβ_^DAI^ localizes to stress granules under conditions of stress (top panels, indicated by arrows). Absence of stress leads to diffuse localization of the protein in both cytoplasm and nucleus (bottom panels). The nucleus is stained with DAPI (blue). TIAR (red) is used as a marker protein for stress granules. **(B)** Immunolocalization of mutant GFP-Z_αβ_^DAI^ (N46A/Y50A/N141A/Y145A) fusion in the presence (+) and absence (−) of stress induced by arsenite in A549 cells. Mutant GFP-Z_αβ_^DAI^ loses the ability to significantly colocalize with TIAR in the stress granules under conditions of arsenite induced stress. The individual panels and the merged images were generated with ImageJ version 1.50i.

### Interaction partners of Z_αβ_^DAI^ and Z_αβ_^DAI^ quadruple mutant under conditions of stress

Triplicate IPs were performed from cells expressing: GFP-Z_αβ_^DAI^, a GFP fusion to a quadruple Z_αβ_^DAI^ mutant that abolishes nucleic acids binding and GFP alone. Each IP was performed with arsenite treatment and all samples were then subjected to LC/MS analysis. Z_αβ_^DAI^ and quadruple Z_αβ_^DAI^ mutant interactome was normalized with respect to GFP alone control. Fig 2A shows the volcano plot depicting the differentially significant interactome for the wild-type Z_αβ_^DAI^ vis-à-vis GFP and Fig 2B shows the same analysis for the quadruple Z_αβ_^DAI^ mutant. Table S2 and S3 summarizes the details schematically presented in Fig 2. The top significant interactors are all found to be RNA binding proteins. The three strongest hits were tRNA-splicing ligase RtcB homolog, 40S ribosomal protein S19 and Nucleolin. These proteins interact independently of whether the nucleic acid binding ability was mutated, suggesting a direct protein interaction. Other proteins that were found as potential interacting partners of both the wild-type and mutant protein were those that are involved in translation (Ribosomal subunits), ATP-dependent RNA helicase activity (DDX5, DDX1, DDX17), mRNA transport (Ras GTPase-activating protein-binding protein 2) and RNA-binding proteins like p54nrb/NONO, SFPQ, RNA-binding protein 14, Nuclear fragile X mental retardation-interacting protein 2, Protein HEXIM1, Heterogeneous nuclear ribonucleoprotein U-like protein 1, Ataxin-2-like protein, Alpha-adducin and FUS. Interestingly, though p54nrb/NONO and FUS are known component of nuclear paraspeckles (Fox and Lamond, 2010), they have also been shown to localize to neuronal cytoplasmic RNP granules (Furukawa et al., 2015). Further, we found the interactions with the small 40S subunit (Z_αβ_^DAI^: S2, S4, S18, S19; Z_αβ_^DAI^ mutant: S2, S5, S6, S11, S14, S18, S19) of ribosome was slightly overrepresented as compared to the large 60S subunit (Z_αβ_^DAI^: P0, 1; Z_αβ_^DAI^ mutant: L26, P0, L19, L8, P1). It has been previously demonstrated that small, but not large, ribosomal subunits are preferably recruited to SGs as shown by immunofluorescent studies with TIAR as the SG marker (Kedersha et al., 2002). Enrichment of ribosomal proteins also suggests that RNAs bound by Z_αβ_^DAI^ were translationally engaged and thus, were potentially associated with ribosomes. Another plausible explanation for the presence of ribosomal proteins could be the demonstrated affinity of Z_α_ domain of ADAR1 for ribosomal RNA segments (Feng et al., 2011).

**Figure 2.**
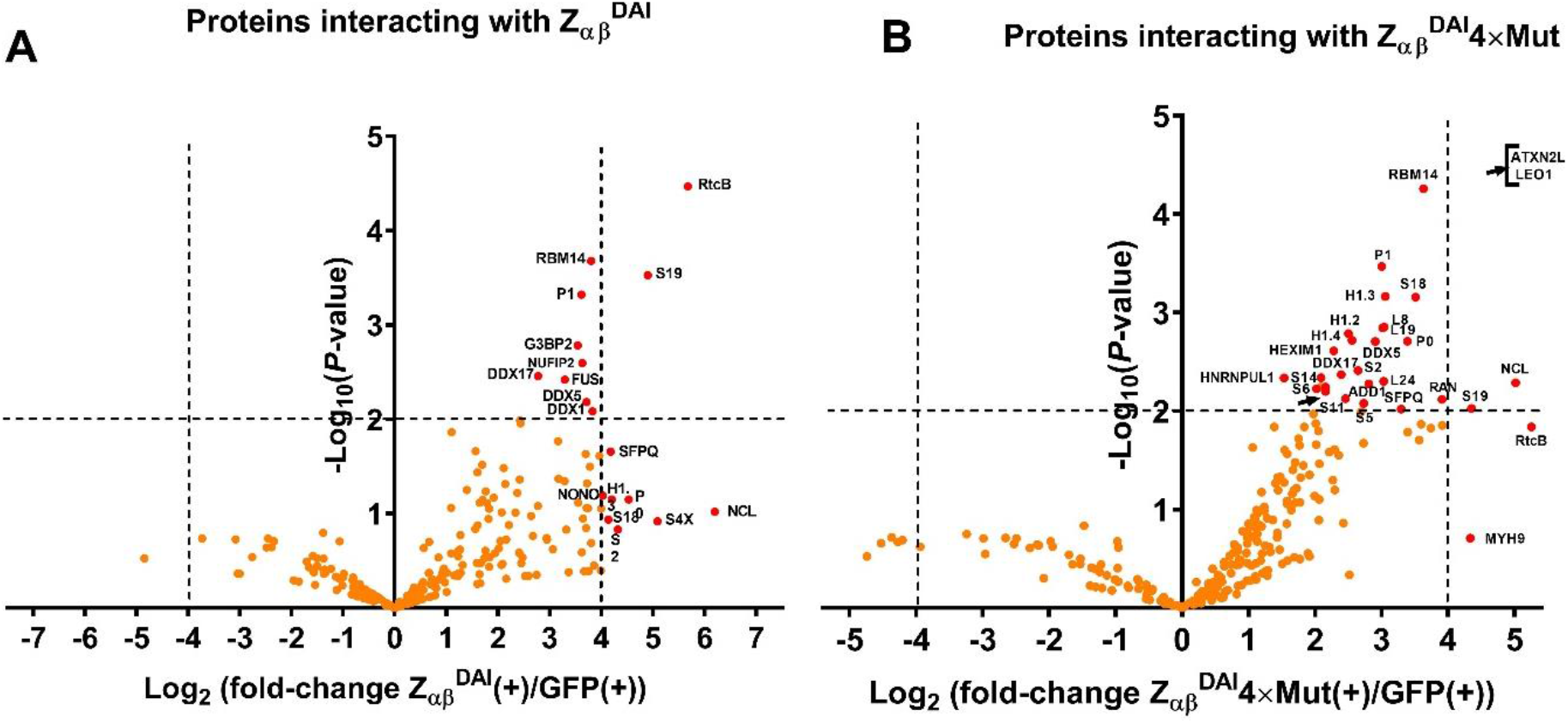
Interaction partners of Z_αβ_^DAI^ protein, normalized with respect to GFP alone control, under conditions of stress induced by arsenite. **A)** Z_αβ_^DAI^ vis-à-vis GFP. **B)** Z_αβ_^DAI^ quadruple mutant vis-à-vis GFP. The proteomics data is plotted as a volcano plot with the –logarithm of p-values computed from a pairwise t-test for each individual interacting protein on the y-axis and the logarithm to the base 2 of the observed fold change for a particular partner with respect to GFP alone control. Data in the center and at the base of the plot represent ~1 fold change and a p-value approaching 1 indicating no significant change. Whereas, points in the upper right and upper left quadrants indicate significant negative and positive changes with respect to the controls. The thresholds (dotted lines) indicate P-value cut-off of 0.01 and a highly stringent fold-change of 16. Dots coloured red are those that are significant with respect to the above mentioned thresholds. The statistical analysis and the plots were generated with the help of GraphPad Prism version 7.02.

### Predominant interactions shown by Z_αβ_^DAI^ with components of SG is stress-independent

To understand whether the induction of stress or its absence significantly changes the nature of Z_αβ_^DAI^ interacting proteins, IPs were performed with Z_αβ_^DAI^ and Z_αβ_^DAI^ quadruple mutant in the presence and absence of stress induced by arsenite and the resultant protein interactants were analysed by LC-MS. Fig 3A and 3B shows the volcano plot for Z_αβ_^DAI^ and Z_αβ_^DAI^ quadruple mutant in the presence and absence of arsenite induced stress, respectively, and Table S4 summarizes the results for the test/control pairs. This analysis reveals no major difference when comparing the interactome in the absence and presence of stress induced by arsenite. This indicates that arsenite-induced stress is not a major driver for selective protein binding to the Z_αβ_^DAI^ domain. Though there are a few proteins that show statistically significant association with respect to their control pair, none of the hits showed both statistical and numerical significance.

**Figure 3.**
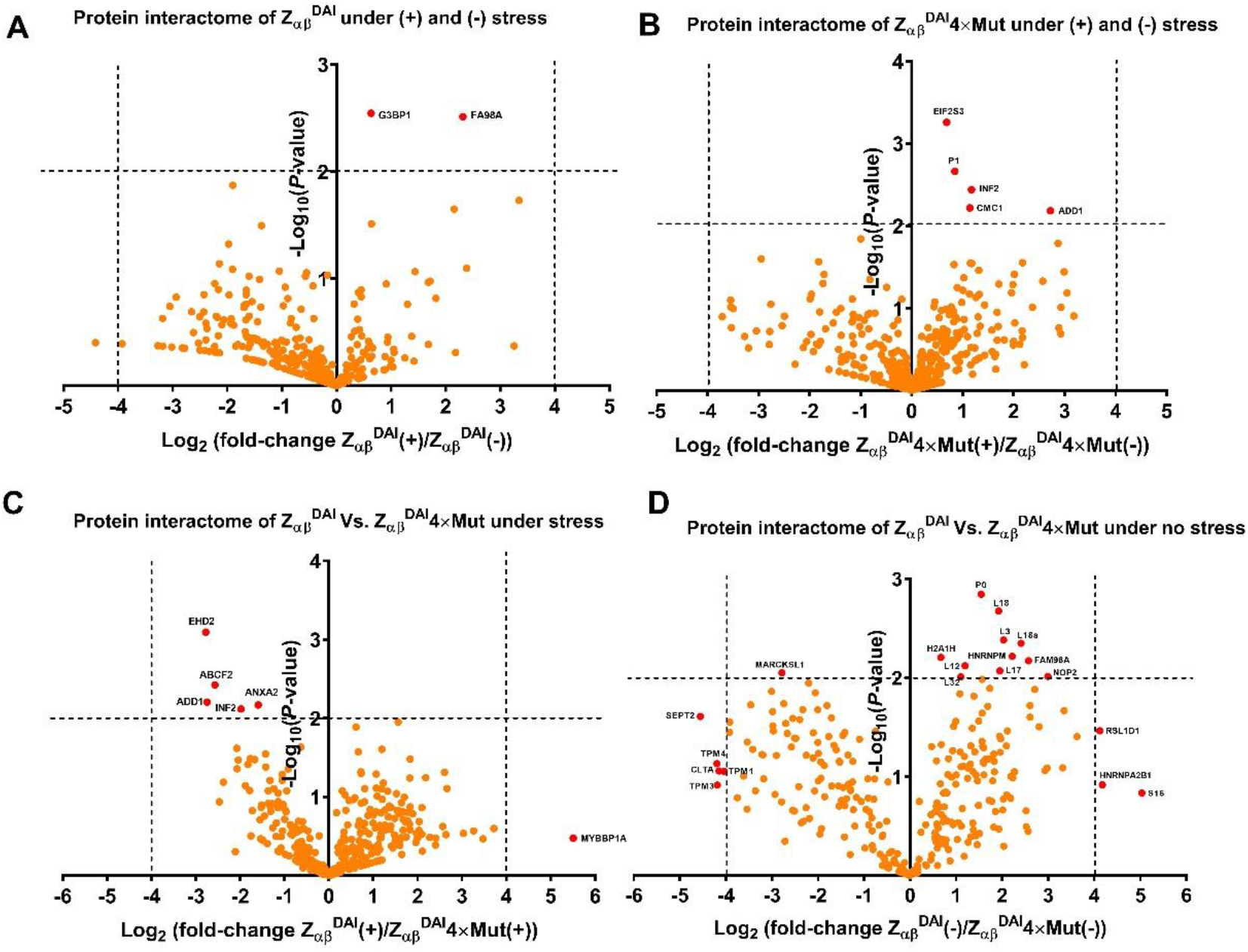
Differential interactions with cellular proteins in the presence and absence of arsenite-induced stress with **(A)** Z_αβ_^DAI^ **(B)** Mutant Z_αβ_^DAI^. Volcano plots on panels (**A**-**B**) provide specific insights into stress granule mediated interactions. Differential interactions with cellular proteins shown by the wild-type and mutant Z_αβ_^DAI^ in the **(C)** presence or **(D)** absence of stress induced by arsenite. Volcano plots on panels (**C**-**D**) provide insights into the interaction enrichment facilitated by the nucleic acid binding ability of the proteins. The volcano plots were plotted as specified in the “materials and methods” section and the thresholds identifying the significance are as discussed in the legend to Fig 2. The statistical analysis and the plots were generated with the help of GraphPad Prism version 7.02.

However, it is noteworthy that the wildtype protein shows preferential association with Ras GTPase-activating protein binding protein 1(G3BP1), a stress granule effector that can induce SG assembly upon overexpression (Tourrière et al., 2003) and FAM98A protein, an RNA-binding protein that has been found in complex with an ATP-dependent RNA helicase DDX1 and C14orf166 (Akter et al., 2017; Pérez-González et al., 2014).

The mutant protein, on the other hand, shows preferential interaction with Eukaryotic translation initiation factor 2 subunit 3, 60S acidic ribosomal protein P1, Inverted formin-2, COX assembly mitochondrial protein homolog, Alpha-adducin under conditions of stress induced by arsenite.

These findings likely reflect the fact that the Z_αβ_^DAI^ that accumulates in SGs represents only a small fraction of the total pool and hence, predominant interactions displayed by them towards other proteins are stress-independent. Thus, it might be reasonable to assume that Z_αβ_^DAI^ may be pre-bound to diffuse mRNA/ribosome associated complexes which may translocate to SGs upon stress induction. Methodologies to isolate SGs specifically are needed to clarify if there are additional partners specific to the stress granule environment. Further, the reported partners are likely specific interactors given that TIAR, a key component of SG, is completely absent in the proteomics data obtained from IPs. This also suggests that the complexes formed by Z_αβ_^DAI^ are diffusely distributed and are likely distinct from the corresponding RNPs.

### Interaction of Z_αβ_^DAI^ domain with other proteins is principally mediated by their nucleic acid binding ability

Comparison of the interacting proteins between wild-type Z_αβ_^DAI^ and the mutant that lacks the ability to bind nucleic acid shows that the bulk of the interactions with ribosomal proteins are mediated by nucleic acid binding ability of the Z_αβ_^DAI^ domain (Fig 3 C and D) (Table S5). Though the results under conditions of arsenite-induced stress are less clear, analysis of the interaction profile of the wild-type and mutant Z_αβ_^DAI^ under physiological condition (no stress) reveals a set of proteins interacting with Z_αβ_^DAI^ in a manner dependent on its nucleic acid binding capacity (Fig 3 D). Under conditions of no arsenite stress, the wild-type protein shows statistically significant difference in interaction with 14 proteins all of which are nucleic acid binding. These proteins include those that constitute 60S and 40S ribosomal subunits, histones and heterogenous nuclear ribonucleoproteins. In complete contrast, the mutant protein shows statistically significant difference in interaction with 6 proteins none of which have been implicated in functions related to RNA binding (Table S5). It should be emphasized though that assigning a cut-off of 2-fold enhancement (log_2_ fold-enhancement =1) at the current *P*-value threshold still follows the patterns whereby all the interactions shown by the wild-type Z_αβ_^DAI^ domain are proteins with nucleic acid binding ability while most interacting partners of the mutant, which have significant statistical and numerical differences than the wild-type, are non-nucleic acid binding proteins (Table S6). This finding unambiguously points to the importance of Z_αβ_^DAI^’s nucleic acid binding ability as the principal driving force behind its interaction with other nucleic acid binding proteins.

### Confirmation of ZαβDAI’s interaction with SG components FUS and p54nrb by Immuno-pulldown and immunofluorescence

To confirm the interaction of Z_αβ_^DAI^ protein with components of SGs as revealed by our MS analysis, we selected two prominent SG associated proteins, FUS and p54nrb. Attempts to probe for SFPQ/PSF were not undertaken as this protein is highly similar to p54nrb (antibodies cross-react) and forms heterodimers with the latter (Yarosh et al., 2015). Western blots on IPs were performed under the same conditions used for the MS characterization. Both FUS and p54nrb were detected in the corresponding IPs (Fig 4). The majority of both proteins was found in the unbound fraction of the IP suggesting that only a fraction of these proteins associate with Z_αβ_^DAI^. This is expected as both proteins have distinct localisation from Z_αβ_^DAI^ and have been primarily reported to be nuclear. To understand if the interaction is dependent on the presence of a nucleic acid intermediate, we performed similar IPs from extracts treated with an RNAse cocktail (Fig 4). Interestingly, P54nrb is lost in RNAse treated samples suggesting an RNA-mediated interaction (Fig 4A). While, FUS levels are significantly reduced but we can still detect the protein present in the IP, pointing to a protein-protein interaction with Z_αβ_^DAI^, at least in part. As a control, we preformed IPs from cells expressing either mutant Z_αβ_^DAI^ or GFP-alone. IPs from the mutant protein shows neither FUS nor P54nrb (Fig 4 A and B). The GFP IPs, likewise, show no p54nrb and FUS, suggesting the interaction is specific (Fig 4). The ability of the mutant Z_αβ_^DAI^ to interact with FUS is unexpected and suggests that the nucleic acid binding activity is required for the FUS interaction despite not mediated by RNA. This indicates that a direct protein-protein interaction is, perhaps allosterically, affected by the engineered mutations or that the RNA and protein interaction surfaces overlap.

**Figure 4.**
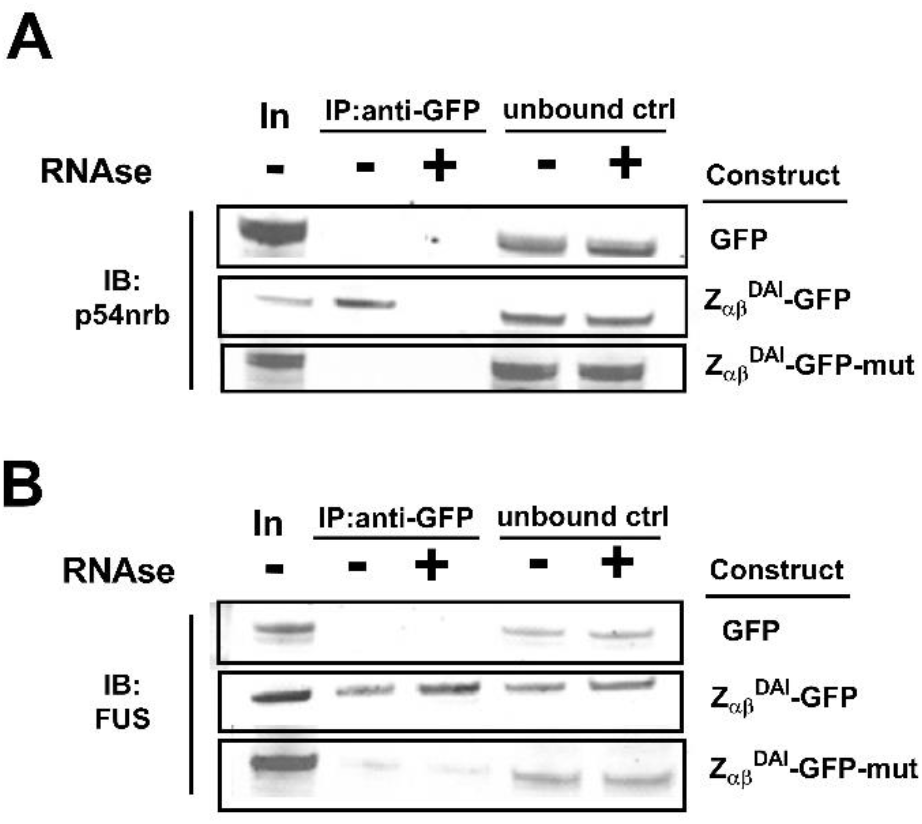
P54nrb forms an RNA dependent complex with Z_αβ_^DAI^. **(A)** Immunoprecipitation of GFP Z_αβ_^DAI^ (middle panel) and mutant GFP Z_αβ_^DAI^ (bottom panel) from the lysate of A549 cells that were immunoblotted with antibodies against p54nrb. The GFP control is shown in the top panel. As seen, p54nrb forms an RNA-dependent complex with Z_αβ_^DAI^ since treatment with RNAse results in complete absence of p54nrb from the blot. **FUS forms an RNA independent complex with Z_αβ_^DAI^. (B)** Immunoprecipitation of GFP Z_αβ_^DAI^ (middle panel) and mutant GFP Z_αβ_^DAI^ (bottom panel) from the lysate of A549 cells that were immunoblotted using antibodies against FUS. The GFP control is shown in the top panel. As seen, FUS forms an RNA-independent complex with Z_αβ_^DAI^ since treatment with RNAse does not change the level of FUS in the blot. However, the results with the mutant are unexpected given that the mutant protein (lacking in vitro nucleic acid binding ability) should have pulled down FUS as much as the wild-type. The abbreviations employed in the figure are I-input, IP-immunoprecipitate, U-unbound and the symbols + and − indicate the presence or absence of RNase. The experiments were performed in triplicate with almost identical outcomes

Next, we aimed to determine where in the cells the Z_αβ_^DAI^/FUS/p54nrb/PSF complex resides. These proteins are known to be localised in the nucleus and have been previously been found enriched in paraspeckles, thus we wondered whether the nuclear speckles observed to accumulate Z_αβ_^DAI^ are paraspeckles. To test this, we co-stained A549 cells with anti-GFP and the antibody for each protein. Neither FUS nor p54nrb are found enriched in Z_αβ_^DAI^ nuclear foci (Fig S2). While, both FUS and p54nrb show a diffuse nuclear staining and consequently we cannot exclude their presence in Z_αβ_^DAI^ speckles, there is no indication of a preferred localisation in these structures. These results suggest that Z_αβ_^DAI^ speckles do not represent paraspeckles. This notion is further supported using PSPC1 staining, another common marker for paraspeckles, which also did not show any specific enrichment in Z_αβ_^DAI^ nuclear speckles (Fig S2).

While we could not detect any co-localization of Z_αβ_^DAI^ with FUS and p54nrb in the nucleus, both proteins were enriched in Z_αβ_^DAI^ foci in the cytoplasm of arsenite treated cells. FUS and p54nrb are enriched in stress granules similarly to what we observe for Z_αβ_^DAI^ (Fig 5). The presence of FUS in stress granules has been described before (Aulas and Velde, 2015) and such presence is known to be enhanced in disease conditions associated with neuro-degenerative disorders (Shelkovnikova et al., 2014). On the other hand, p54nrb and PSF are not commonly found to be present in stress granules, except thus far in mouse retinal cell lines (Furukawa et al., 2015). Thus, our results point to a possible novel p54nrb/PSF function at stress granules.

**Figure 5.**
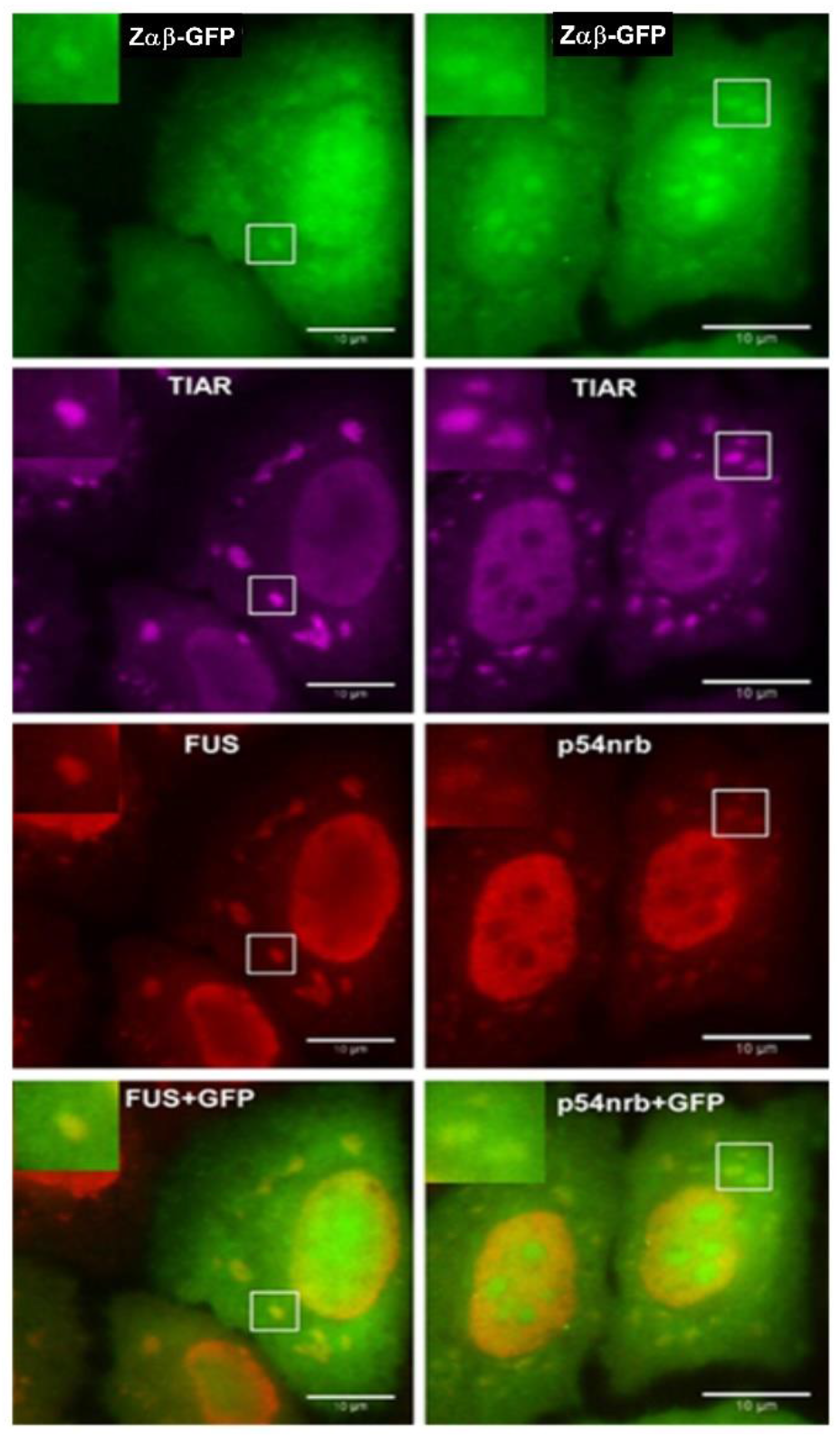
Both p54nrb and FUS along with Z_αβ_^DAI^ are enriched in stress granules. Arsenite treated A549 cells show enrichment of GFP-ZαβDAI (green) in cytoplasmic aggregates that are identified as stress granules by co-staining for TIAR (magenta). The same cytoplasmic bodies are stained by anti-p54nrb and anti-FUS antibodies despite an intense nuclear localisation for both proteins

## Discussion

The ADAR1 Zα domain was discovered in an *in vitro* screen aimed to identify Z-DNA binding proteins. Whether the ability of the domain to do so was linked to its *in vivo* function or coincidental has been a lingering question that is yet not fully clarified. Support for the *in vivo* significance of Zα domain binding to the left-handed helical conformation of nucleic acid came from the discovery of other proteins with Zα domains and the demonstration that all these domains behave similarly *in vitro* and share invariant amino acids shown to be crucial for Z-DNA binding. Crucially, replacement of E3L Zα domains with those of ADAR1 or DAI showed that they can functionally replace each other and mutagenesis in this context has shown that amino acids crucial for Z-DNA binding are also essential for the *in vivo* function.

Although the interaction of Zα with Z-DNA has been heavily studied, it became clear early on that Zα domains can interact with the left-handed RNA double helix as well. Considering the known functions of the proteins that have Zα domains, an RNA targeting function appears more likely. The results presented here further support an interaction of Zα domains with RNA. The enrichment of Zα domains in stress granules, a location of storage for stalled ribosomes and mRNAs, suggests that Zα domains are pre-bound to mRNAs or ribosomal RNA which translocate to SGs resulting in the observed enrichment. It has been suggested that Zα domains are specifically targeted to SGs, however our results do not support this notion as only a fraction of Zα is localised to SGs and rather suggest a passive increase due to RNA enrichment. Indeed, with a few notable exceptions, we do not observe major differences in proteins that interact with the Zα domain and RNA in the presence or absence of stress.

All the top interacting proteins we identify are previously known RNA interacting proteins with key roles in RNA processing and trafficking. Among them p54nrb or NONO is of particular interest as it has been previously indirectly linked to ADARs and Zα by showing to be involved in nuclear retention of heavily edited mRNAs (Prasanth et al., 2005). It has been speculated that p54nrb may directly recognise the presence of inosine in such mRNAs but no direct evidence of this recognition has emerged yet. This previously described p54nrb interaction with edited transcripts is nuclear and linked to the formation of paraspeckles. We found that p54nrb is also present in the cytoplasm albeit at much lower levels. However, significant amounts of the protein are found in stress granules. Whether this fraction of p54nrb originates from a diffuse cytoplasmic distribution or directly migrates from the nucleus is not clear and needs further study. Our experiments suggest that the nuclear Z_αβ_^DAI^ speckles are not enriched in p54nrb and are not paraspeckles and thus this is not likely were p54nrb and Z_αβ_^DAI^ complex originates. The identification of SFPQ as a top interactor is not surprising given the presence of p54nrb, the two proteins are homologues and known to form heterodimers consistent with our finding of similar number of peptides for the two proteins.

FUS shows a similar localization distribution as p54nrb but its translocation to stress granules has been previously observed and linked to pathologies leading to neurodegenerative disorders (Li et al., 2013; Masuda et al., 2016). Thus, an interpretation of our results would be that Zα domains target a subset of mRNAs also targeted by a FUS/p54nrb/SFPQ complex. The function of such a complex and the nature of RNAs targeted are open questions and will be discussed further. Interestingly FUS mutations cause Amyotrophic Lateral Sclerosis (ALS) that leads to cell death of motor neurons (An et al., 2019). Cell death in this disease has been attributed to hypoediting of GLUR2 subunit of AMPA receptors leading to a lethal increase of intracellular calcium (Da Cruz and Cleveland, 2011). How FUS mutations are linked to changes in editing levels has been unclear. Although GLUR2 editing is mediated by ADAR2 (Peng et al., 2006), changes in ADAR1, which is known to compete with ADAR2 for substrate binding, may influence ADAR2 activity. This way, our findings provide the first link between FUS and the editing machinery that might be worth exploring further in the context of ALS. Several other RNA binding proteins (DDX1, HNRNPUL1, RTCB, and C14orf166) are also found to be enriched in Z_αβ_^DAI^ IPs. This includes several helicases and splicing related proteins, supporting the idea that the entire complex accompanies trafficking mRNAs.

A significant effort has been devoted to characterising the RNA binding properties of FUS using CLIP-seq and *in vitro* SELEX analysis, revealing a complex RNA recognition pattern (Wang et al., 2015) (Choi et al., 2017). CLIP-seq experiments show a recognition of GU-rich motifs and similarly SELEX revealed GGUG motif recognition with RNA recognition shown to be direct and mediated by the zinc-finger domain of FUS (Loughlin et al., 2019). Other studies, however, suggest limited specificity and indicate that GU motifs cannot explain the range of RNA binding interactions of FUS (Wang et al., 2015). RNA binding close to alternative splicing signals and regulation of alternative splicing is the most prevalent functional role assigned to FUS although links to transcription and polyadenylation have been reported (Masuda et al., 2015). Given the multifunctional role of FUS and the lack of clear specificity, it is possible that specific functions of FUS are mediated by other proteins providing sequence or structural specificity. Our data suggest that the interaction of Z_αβ_^DAI^ with FUS includes a protein-protein component and thus it is tempting to suggest that interactions with Zα domains may guide the cytoplasmic functions of FUS.

PSF (PTB-associated splicing factor) and NONO/p54nrb along with PSPC1 form the DHBS family of proteins which share homology, domains and a multifunctional role centred on splicing and mRNA metabolism (Yarosh et al., 2015). In the nucleus PSF and NONO are found in paraspeckles where they interact with the scaffolding lncRNA Neat1 that forms the basis of paraspeckles. The two proteins are shown to mediate nuclear retention of heavily edited RNAs by ADARs and thus their presence as strong interactors with Zα domains are of particular interest. However, we observe no colocalization of Z_αβ_^DAI^ with PSF/NONO in nuclear structures suggesting that the signal we detect may come from cytoplasmic complexes. While PSF and NONO are primarily nuclear, reports show a role of PSF in IRES mediated translation (Sharathchandra et al., 2012) and translocation of PSF in the cytoplasm in Alzheimer’s and Pick’s disease (Ke et al., 2012). Our results showing NONO localising in arsenite induced stress granules emphasize a cytoplasmic role for PSF/NONO.

What is the nature of the RNAs on which Z_αβ_^DAI^ and FUS/NONO/PSF are bound? Given the described affinity of NONO/PSF for inosine containing duplexes, it is tempting to speculate that edited RNAs that are not retained in (or escape from) paraspeckles are targeted in the cytoplasm and escorted to SGs upon elevated stress. Since it is known that ADAR1 Zα targets RNAs that have already been modified in the nucleus, selective targeting of nucleic acid by the enzyme for more extensive modification in the cytoplasm is highly likely. Although the propensity and extent of inosine containing RNA duplexes to be in the left-handed conformation is not clear, it is tempting to suggest that selective binding of Zα to the left-handed helical conformation of I-containing nucleotides targets proteins to modified duplexes without the need to specifically recognise the inosine base. We now know that the long cytoplasmic form of ADAR1 which contains the Zα domain has a crucial role in restoring homeostasis after the activation of the IFN pathway. Targeting of ADAR1 specifically on activating RNAs would efficiently reduce cytoplasmic danger signals while avoiding binding and sequestration of the enzyme on innocuous RNAs.

## Abbreviations

ADAR1: Adenosine deaminases acting on RNA
SG: Stress granules
NONO: Non-POU domain-containing octamer-binding protein
SFPQ: splicing factor proline and glutamine rich
G3BP1: Ras GTPase-activating protein-binding protein 1
DDX5: DEAD-Box Helicase 5
DDX1: DEAD-Box Helicase 1
RBM14: RNA Binding motif Protein 14
LC-MS: Liquid chromatography mass spectrometry
DAI: DNA-dependent activator of IFN-regulatory factor

## Dedication

This manuscript is dedicated to the memory of Alekos Athanasiadis who sadly passed away in August 2020. Though the manuscript is communicated by BS for practical purposes, the work was conceived and supervised by AA.

## Acknowledgements

We thank Élio Sucena, Jonathan C. Howard and Mónica Bettencourt-Dias for providing useful comments. This work was supported by a Fundação para a Ciência e Tecnologia (FCT) grant PTDC/BBB-BEP/3380/2014 to LG, BS and AA. BS was also supported by Marie Skłodowska-Curie Individual Fellowship Grant 789565. KK was a recipient of FCT PhD grant SFRH/BD/51626/2011. MJA was supported by the grant PTDC/BIA-CEL/32211/2017 awarded by the FCT, Portugal. LETJ was supported by an EMBO Installation Grant 1818, an ERC-consolidator grant ERC-2013-CoG-615638 and an “Investigador FCT” position.

