## Supplemental material for "Enrichment of Zα domains at cytoplasmic stress granules is due to their innate ability to bind nucleic acids"

### Results:

#### $Z_{\alpha\beta}^{DAI}$ foci represent a novel nuclear structure

Apart from cytoplasmic stress granules, $Z_{\alpha\beta}^{DAI}$ also localizes to nuclear speckles (Fig S1). However, the physiological significance of such localization is not clear since none of the $Z_{\alpha\beta}^{DAI}$ containing proteins are localized in the nucleus. It is likely that the $Z_{\alpha\beta}^{DAI}$ protein localizes to the nucleolus because nucleolus contains high concentrations of ribosomal RNAs and $Z_{\alpha\beta}^{DAI}$ binds ribosomal RNA with high affinity (Feng et al., 2011). However, we decided to pursue this angle of research since, $Z_{\alpha\beta}^{DAI}$ can be utilized as a novel nuclear marker depending on its localization to, possibly, unique nuclear substructures. To understand the nuclear localization of $Z_{\alpha\beta}^{DAI}$, colocalization of the protein was attempted with paraspeckle markers P54nrb and FUS. $Z_{\alpha\beta}^{DAI}$ failed to colocalize with the marker proteins in the nucleus (Fig S2). Hence, it can be concluded that nuclear structures enriched in $Z_{\alpha\beta}^{DAI}$ do not represent paraspeckles. We decided to examine if they coincide with another known nuclear structure such as PML and Cajal bodies and whose general properties are consistent with the ones observed for $Z_{\alpha\beta}^{DAI}$ speckles. However, co-staining of arsenite treated cells with either anti-PML or anti-coilin antibodies shows that, despite their close resemblance, $Z_{\alpha\beta}^{DAI}$ bodies are indeed distinct structures (Figure S3). This nuclear accumulation of $Z_{\alpha\beta}^{DAI}$ in dynamically formed bodies that do not coincide with paraspeckles, speckles or PML bodies is a puzzling feature of Zα domains described here. To our knowledge, $Z_{\alpha\beta}^{DAI}$ bodies are the first nuclear structure whose formation is induced by stress. Although other structures such as PML bodies are responsive to stress, such bodies' existence is not stress dependent. Given that the presumed function of Zα domains appears to be in the cytoplasm, it is not clear that the accumulation of $Z_{\alpha\beta}^{DAI}$ in these structures is physiologically relevant. Nevertheless, Zα domains may prove a valuable tool to characterise a novel nuclear structure of relevance to immune and stress responses.

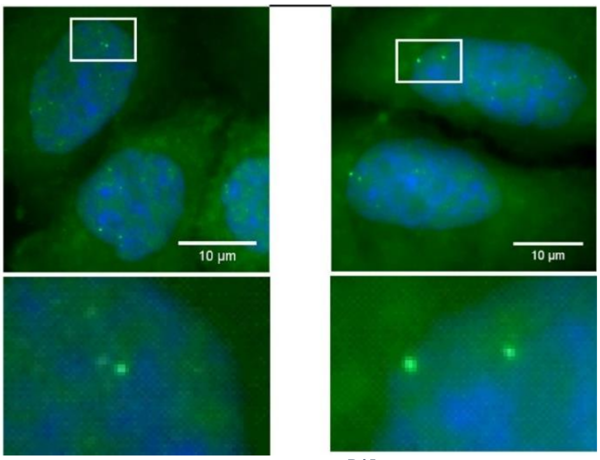


***Figure S1. Accumulation of $Z_{\alpha\beta}^{DAI}$ in the nuclear bodies.*** *$Z_{\alpha\beta}^{DAI}$ is also enriched in nuclear bodies that resemble paraspeckles. Arsenite treated cells show an accumulation of $Z_{\alpha\beta}^{DAI}$ in dot-like nuclear bodies. Nuclear $Z_{\alpha\beta}^{DAI}$ bodies vary in number between cells and on average there are 10 per cell.The figure shows two examples of the same condition.*

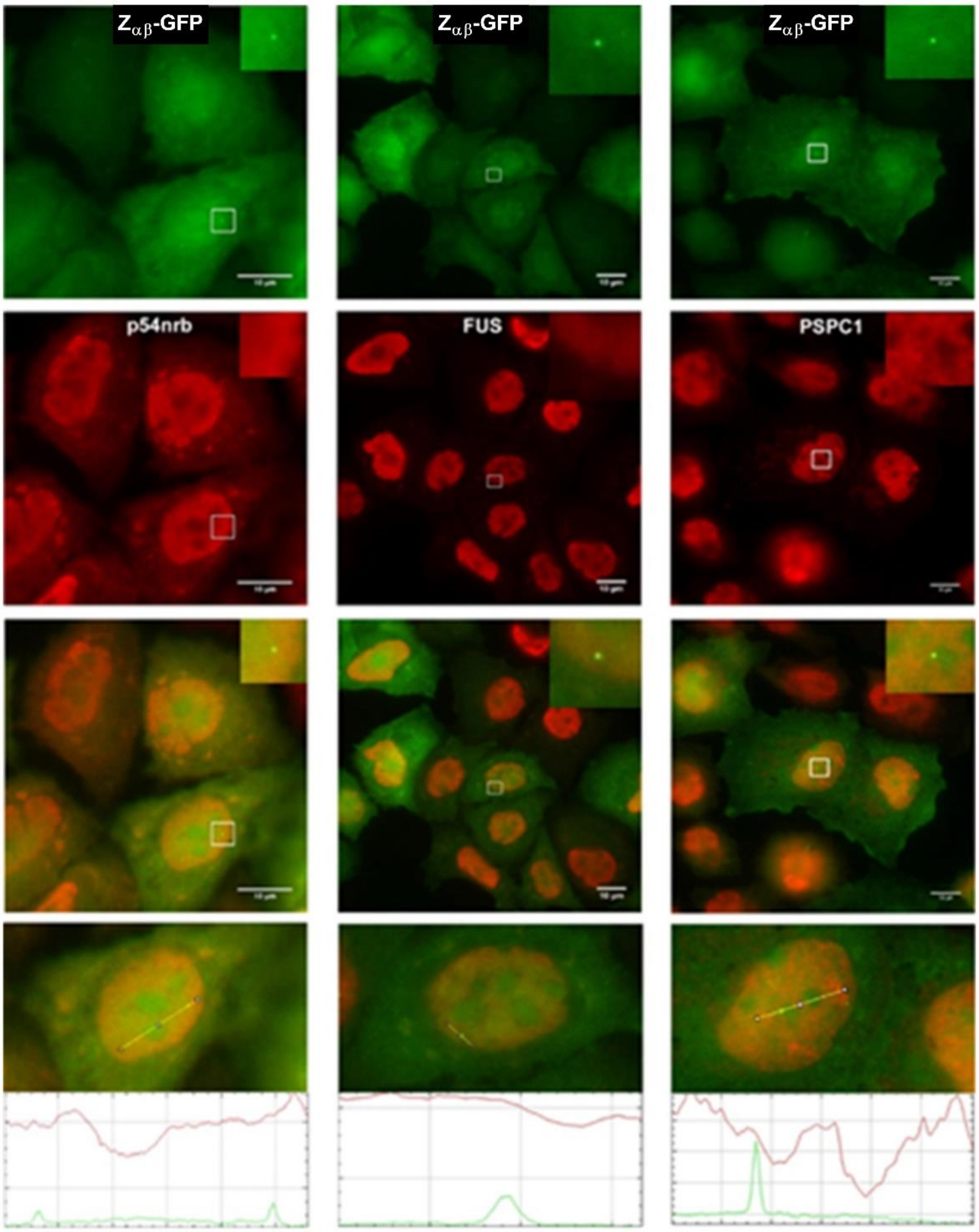


***Figure S2. P54nrb and FUS are notspecificallyenriched in $Z_{\alpha\beta}^{DAI}$ positive nuclear speckles.*** *Insets at the upper right of each photograph are magnifications of the square encompassing a single $Z_{\alpha\beta}^{DAI}$ nuclear body. PSPC1 is a core protein of paraspeckles absent in the $Z_{\alpha\beta}^{DAI}$ IPs used as a control. Enrichment of $Z_{\alpha\beta}^{DAI}$ in nucleolus is in contrast to its avoidance by p54nrb and FUS. Bottom row shows quantification along the indicated lines (Green $Z_{\alpha\beta}^{DAI}$; red the corresponding protein).*

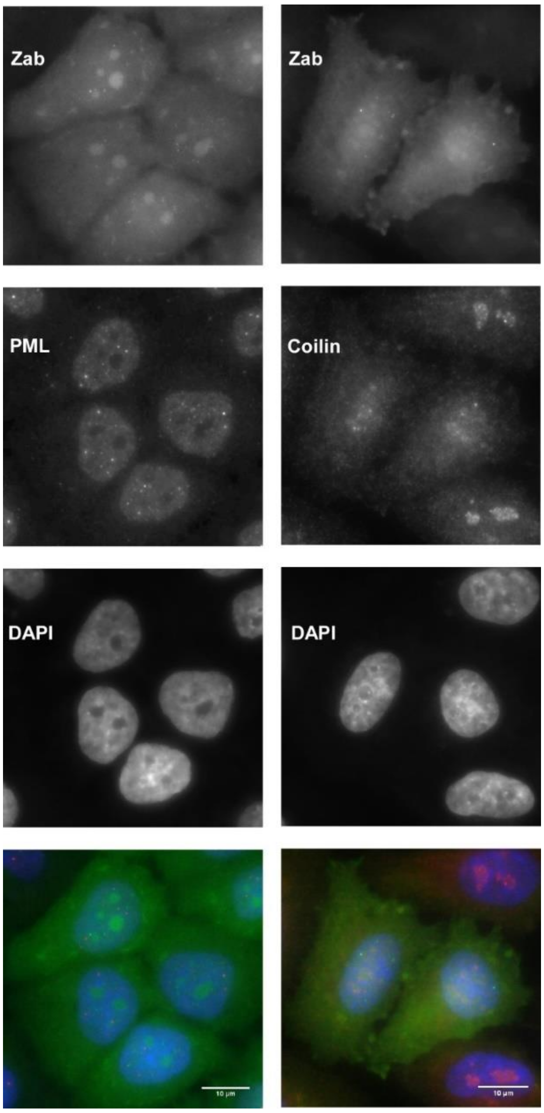


***Figure S3. $Z_{\alpha\beta}^{DAI}$ nuclear aggregates do not coincide with PML and Cajal bodies****. A549 cells treated with arsenite are visualised for GFP- $Z_{\alpha\beta}^{DAI}$ (green) and co-stained for PML or coilin. DAPI staining is in blue to indicate nuclear boundaries*

### Supplementary Tables

**Table S1**. Primers used for Zαβ expression constructs.

| Reference | Primer name | Sequence |
|---|---|---|
| 1 | ForZabhDAI | ATC GAT CTC GAG GCC CAG GCT CCT GCT GAC CCG |
| 2 | RevZabhDAI | GCG CGC GGA TCC TTA GTA AAT CGT CCA TGC TTT GGA CTG |
| 3 | ForGFPBamHI | ATC GAT GGA TCC ATG GTG AGC AAG GGC GAG GAG |
| 4 | RevZbDAISalI | GCG CGC GTC GAC TTA GTA AAT CGT CCA TGC TTT GGA |
| 5 | RevSprot | GCG CGC GTC GAC TTA ACT AGT ACC TCC ACC TCC GCT |
| 6 | ForZaDAIMut | CCC AAG AGG GAG CTC GCG CAA GTC CTC GCG CGA ATG AAA AAG GAG TTG |
| 7 | RevZaDAIMut | CAA CTC CTT TTT CAT TCG CGC GAG GAC TTG CGC GAG CTC CCT CTT GGG |
| 8 | ForZbDAIMut | ACA GCA AAA GAT GTG GCG CGA GAC TTG GCG AGG ATG AAG AGC AGG CAC |
| 9 | RevZbDAIMut | GTG CCT GCT CTT CAT CCT CGC CAA GTC TCG CGC CAC ATC TTT TGC TGT |

**Table S2.** Proteins that show significant enrichment in wild-type $Z_{\alpha\beta}^{DAI}$ treated with arsenite vis-à-vis treated GFP samples as assessed by volcano plot that weights the significance based on the dual criterion of numerical fold-increase and statistical p-value significance. The gene

| Significance | Uniprot ID | Description |
|---|---|---|
| Both p-value and fold-change | Q9Y3I0 | tRNA-splicing ligase RtcB homolog(RtcB) |
| | P39019 | 40S ribosomal protein S19 (S19) |
| Fold-change | P19338 | Nucleolin (NCL) |
| | P62701 | 40S ribosomal protein S4, isoform X (S4X) |
| | P05388 | 60S acidic ribosomal protein P0 (P0) |
| | P15880 | 40S ribosomal protein S2 (S2) |
| | P62269 | 40S ribosomal protein S18 (S18) |
| | P16402 | Histone H1.3 (H1.3) |
| | Q15233 | Non-POU domain-containing octamer-binding protein (NONO) |
| | P23246 | Splicing factor, proline- and glutamine-rich (SFPQ) |
| P-value significance | Q96PK6 | RNA-binding protein 14 (RBM14) |
| | P05386 | 60S acidic ribosomal protein 1 |
| | Q9UN86 | Ras GTPase-activating protein-binding protein 2 (G3BP2) |
| | Q92841 | Probable ATP-dependent RNA helicase DDX17 (DDX17) |
| | Q7Z417 | Nuclear fragile X mental retardation-interacting protein 2 (NUFIP2) |
| | P35637 | RNA-binding protein FUS (FUS) |
| | P17844 | Probable ATP-dependent RNA helicase DDX5 (DDX5) |
| | Q92499 | ATP-dependent RNA helicase DDX1 |

name abbreviations in parenthesis indicate the usage in Figure 2.

**Table S3.** Proteins that show significant enrichment in mutant $Z_{\alpha\beta}^{DAI}$ treated with arsenite vis-à-vis treated GFP samples as assessed by volcano plot that weights the significance based on the dual criterion of numerical fold-increase and statistical p-value significance. The gene name abbreviations in parenthesis indicate the usage in Figure 2.

| Significance | Uniprot ID | Description |
|---|---|---|
| Both p-value and Fold-change | P19338 | Nucleolin (NCL) |
| | P39019 | 40S ribosomal protein S19 (S19) |
| | Q9Y3I0 | tRNA-splicing ligase RtcB homolog (RtcB) |
| | P35579 | Myosin 9 (MYH-9) |
| P-value significance | Q96PK6 | RNA-binding protein 14 (RBM14) |
| | P62826 | GTP-binding nuclear protein Ran (RAN) |
| | P05386 | 60S acidic ribosomal protein P1 (P1) |
| | P62269 | 40S ribosomal protein S18 (S18) |
| | P16402 | Histone H1.3 (H1.3) |
| | P62917 | 60S ribosomal protein L8 (L8) |
| | P84098 | 60S ribosomal protein L19 (L19) |
| | P05388 | 60S acidic ribosomal protein P0 (P0) |
| | P17844 | Probable ATP-dependent RNA helicase DDX5 (DDX5) |
| | P16403 | Histone H1.2 (H1.2) |
| | P10412 | Histone H1.4 (H1.4) |
| | O94992 | Protein HEXIM1 (HWXIM1) |
| | Q9BUJ2 | Heterogeneous nuclear ribonucleoprotein U-like protein 1 (HNRNPUL1) |
| | P15880 | 40S ribosomal protein S2 (S2) |
| | Q92841 | Probable ATP-dependent RNA helicase DDX17 (DDX17) |
| | P62263 | 40S ribosomal protein S14 (S14) |
| | P61254 | 60S ribosomal protein L26 (L26) |
| | Q8WWM7 | Ataxin-2-like protein (ATXN2L) |
| | P62753 | 40S ribosomal protein S6 (S6) |
| | P62280 | 40S ribosomal protein S11 (S11) |
| | P35611 | Alpha-adducin (ADD1) |
| | P46782 | 40S ribosomal protein S5 (S5) |
| | P23246 | Splicing factor, proline and glutamine rich (SFPQ) |
| | P83731 | 60S ribosomal protein L24 (L24) |
| | Q8WVC0 | RNA-polymerase associated protein LEO 1 (LEO1) |

**Table S4.** Proteins that show significant enrichment upon treatment with arsenite in both wild-type and mutant $Z_{\alpha\beta}^{DAI}$ proteins as assessed by volcano plot indicating towards preferential interacting partners during SG formation, if any. It should be noted that all of the reported hits have significant p-value differences with respect to no stress conditions while displaying fold-changes below the arbitrary threshold of significance. The gene name abbreviations in parenthesis indicate the usage in Figure 3.

| Significance | Uniprot | Description |
|---|---|---|
| $Z_{\alpha\beta}^{DAI}$ wild-type | Q13283 | Ras GTPase-activating protein binding protein 1 (G3BP1) |
| | Q8NCA5 | Protein FAM98A (FA98A) |
| $Z_{\alpha\beta}^{DAI}$ mutant | P41091 | Eukaryotic translation initiation factor 2 subunit 3 (EIF2S3) |
| | P05386 | 60S acidic ribosomal protein P1 (P1) |
| | Q27J81 | Inverted formin-2 (INF2) |
| | Q7Z7K0 | COX assembly mitochondrial protein homolog (CMC1) |
| | P35611 | Alpha-adducin (ADD1) |

**Table S5.** Differential interacting partners for the $Z_{\alpha\beta}^{DAI}$ wild-type and $Z_{\alpha\beta}^{DAI}$ mutant domain (in the presence and absence of arsenite) indicating interactions mediated by nucleic acid or

| Arsenite | Protein | Uniprot | Description |
|---|---|---|---|
| (+) | $Z_{\alpha\beta}^{DAI}$ | Q9BQG | Myb-binding protein 1A (MYBBP1A) |
| | $Z_{\alpha\beta}^{DAI}$ mutant | Q9NZN4 | EH-domain containing protein 2 (EHD2) |
| | | Q9UG63 | ATP-binding cassette sub-family F member 2 (ABCF2) |
| | | Q27J81 | Inverted formin 2 (INF2) |
| | | P35611 | Alpha-adducin (ADD1) |
| | | P07355 | Annexin A2 (ANXA2) |
| Arsenite (-) | $Z_{\alpha\beta}^{DAI}$ | O76021 | Ribosomal L1 domain containing protein (RSL1D1) |
| | | P22626 | Heterogeneous nuclear ribonucleoproteins A2/B1 (HNRNPA2B1) |
| | | P62249 | 40S ribosomal protein S16 (S16) |
| | | P05388 | 60S acidic ribosomal protein P0 (P0) |
| | | Q07020 | 60S ribosomal protein L18 (L18) |
| | | P39023 | 60S ribosomal protein L3 (L3) |
| | | Q02543 | 60S ribosomal protein L18a (L18a) |
| | | Q8NCA | Protein FAM98A (FAM98A) |
| | | P46087 | Probable 28S rRNA (cytosine(4447)-C(5))-methyltransferase (NOP2) |
| | | P52272 | Heterogeneous nuclear ribonucleoprotein M (HNRNPM) |
| | | P18621 | 60S ribosomal protein L17 (L17) |
| | | Q96KK5 | Histone H2A type 1-H (H2A1H) |
| | | P30050 | 60S ribosomal protein L12 (L12) |
| | | P62010 | 60S ribosomal protein L32 (L32) |
| | $Z_{\alpha\beta}^{DAI}$ mutant | P49006 | MARCKS-related protein (MARCKSL1) |
| | | Q15019 | Septin-2 (SEPT2) |
| | | P67936 | Tropomyosin alpha-4 chain (TPM4) |
| | | P09496 | Clathrin light chain A (CLTA) |
| | | P09493 | Tropomyosin alpha-1 chain (TPM1) |
| | | | Tropomyosin alpha-3 chain (TPM3) |

protein interfaces. * is CON_007144053 and the gene name abbreviations in parenthesis indicate the usage in Figure 3.
